# A hybrid computational model of cancer spheroid growth with ribose-induced collagen stiffening

**DOI:** 10.1101/2024.10.23.619655

**Authors:** Margherita Botticelli, John Metzcar, Thomas Phillips, Susan Cox, Pradeep Keshavanarayana, Fabian Spill

## Abstract

Metastasis, the leading cause of death in cancer patients, arises when cancer cells disseminate from a primary solid tumour to distant organs. Growth and invasion of the solid tumour often involve collective cell migration, which is profoundly influenced by cell-cell interactions and the extracellular matrix (ECM). The ECM’s biochemical composition and mechanical properties, such as stiffness, regulate cancer cell behaviour and migration dynamics. Mathematical modelling serves as a pivotal tool for studying and predicting these complex dynamics, with hybrid discrete-continuous models offering a powerful approach by combining agent-based representations of cells with continuum descriptions of the surrounding microenvironment. In this study, we investigate the impact of ECM stiffness, modulated via ribose-induced collagen cross-linking, on cancer spheroid growth and invasion. We employed a hybrid discrete-continuous model implemented in PhysiCell to simulate spheroid dynamics, successfully replicating three-dimensional *in vitro* experiments. The model incorporates detailed representations of cell-cell and cell-ECM interactions, ECM remodelling, and cell proliferation. Our simulations align with experimental observations of two breast cancer cell lines, non-invasive MCF7 and invasive HCC1954, under varying ECM stiffness conditions. The results demonstrate that increased ECM stiffness due to ribose-induced cross-linking inhibits spheroid invasion in invasive cells, whereas non-invasive cells remain largely unaffected. Furthermore, our simulations show that higher ECM degradation by the cells not only enables spheroid growth and invasion but also facilitates the formation of multicellular protrusions. Conversely, increasing the maximum speed that cells can reach due to cell-ECM interactions enhances spheroid growth while promoting single-cell invasion. This hybrid modelling approach enhances our understanding of the interplay between cancer cell migration, proliferation, and ECM mechanical properties, paving the way for future studies incorporating additional ECM characteristics and microenvironmental conditions.

## 1 Introduction

The extracellular matrix (ECM) is a complex network of numerous macromolecules present within all tissues outside the cells. It comprises approximately 300 different proteins, with fibrous proteins such as collagens being the most abundant^1^. The ECM’s composition and structure, which vary based on tissue type and location, determine distinct mechanical and biochemical properties, which regulate tissue homeostasis, cell differentiation and growth, and largely influence cell migration ^2^. The physical and mechanical characteristics of the ECM, including fibre orientation, stiffness, viscoelasticity and porosity, affect the cell’s direction of movement, speed and various modes of migration (single or collective) ^3^. The ECM plays an important role in regulating cell migration during cancer metastasis, which is the primary cause of death in cancer patients^4^. More mesenchymal cancer cell lines typically invade as single cells and this mode of invasion is profoundly impacted by ECM pore and fibre size ^5^. In solid tumours, when cancer cells migrate collectively, their movement is directed by both cell-cell interactions and their interaction with the ECM ^6^. When cells interact with the ECM, they sense and respond to mechanical cues from the ECM through mechanotransduction pathways, often mediated by integrin signalling ^7^. These cues can trigger intracellular signalling cascades, ultimately impacting cell behaviour ^2,8^. The tumour microenvironment differs significantly from healthy microenvironments and is heavily remodelled by cancer cells and fibroblasts ^9^. Understanding how cancer cells sense and respond to the ECM and its mechanical and biochemical features is crucial to better understanding cancer cell migration and invasion.

Over the years, researchers have investigated cell-ECM interactions in relation to various mechanical properties of the ECM, including stiffness, a material’s resistance to deformation. Notably, increases in ECM stiffness can happen during tumour progression and affect cancer cell behaviour and migration ^10–12^. Furthermore, higher ECM stiffness in breast cancers is usually associated with poor prognosis and drug resistance ^12,13^. Many studies looking at ECM stiffness employ *in vitro* models that simplify the ECM composition, for example by using only collagen to represent the ECM, and are performed in two dimensions due to ease of sample generation, their simplicity in analysing the results and easier reproducibility ^11^. However, there are intrinsic differences between cell cultures in two-dimensional (2D) and three-dimensional (3D) matrices. Cell migration features, such as migration speed, directionality, cell morphology and cytoskeletal organisation are profoundly influenced by the surrounding ECM, while key factors seen *in vivo* such as ECM remodelling inherently require an ECM component^14,15^. For example, cells on 2D matrices present a flatter and more spread-out morphology with large flat protrusions (lamellipodia) mainly localised at the leading edge of the cell. In 3D the cells have a more varied morphology and form protrusions on the whole cell surface, such as pseudopodia and invadopodia, adapting to the surrounding ECM ^16^. Therefore, to faithfully replicate 3D *in vivo* tumour microenvironments, we must focus on 3D *in vitro* ECM models rather than relying solely on 2D models. However, creating 3D *in vitro* ECM models presents challenges due to their complexity, both in sample preparation and data analysis, which explains why researchers often turn to 2D experiments to study mechanical effects of the ECM.

A recent review by Micalet et al. ^11^ collected various papers investigating the effects of ECM stiffness in 3D *in vitro* models of epithelial cancer cells. Some studies found that matrices with higher stiffness can enhance cell migration, and promote epithelial-to-mesenchymal transition (EMT), a process associated with increased invasiveness ^17,18^. On the other hand, other studies have found that less stiff matrices can drive more invasive phenotypes in cancer cells and spheroids ^12,19,20^. Thus, the effect of ECM on cancer migration and invasion depends not only on stiffness, but on several factors, including the cancer cell type, the involvement of other cells such as fibroblasts, and the composition and structure of the surrounding microenvironment. Furthermore, ECM stiffness in *in vitro* models can be modulated using different techniques, each affecting cell migration differently. One approach involves increasing the collagen density and therefore the extracellular matrix stiffness. However, this alters the structural properties of the ECM, such as its porosity, and influences cell behaviour, including the formation and number of focal adhesions (cell-ECM adhesion sites), which often alters cell migration ^21^. Another method is to modify the alginate hydrogel density. However, this limits cell migration by preventing chemical remodelling of the ECM, essential for cell invasion ^21^. In other studies, the stiffness of the collagen matrix is modulated using non-enzymatic glycation, induced by sugars such as threose and ribose. This process increases cross-links among collagen fibres, increasing fibre stiffness without modifying the structure of the ECM^12,19^. The different approaches used in modelling and measuring cancer cell invasion in 3D lead to contradictory results which are also difficult to compare. It is therefore even harder to understand the mechanisms behind cancer cell invasion and their dependence on ECM stiffness.

Given the complexity, cost, and duration of *in vitro* experiments, computational models have become valuable tools in complementing experimental work by replicating setups, providing insights that help interpret results, and exploring scenarios that are difficult to test experimentally^22,23^. Cancer spheroids, commonly used in 3D *in vitro* models to investigate tumour growth and invasion, frequently display collective invasion behaviour. Various computational approaches are employed to model this collective migration, including continuous, discrete, and hybrid models, all of which have been used to examine cancer invasion and interactions between the cells and the surrounding extracellular matrix. In continuous models, the cells and the ECM are represented as densities, characterising them with partial differential equations to describe how they change in space and time ^24,25^. In discrete frameworks, agent-based modelling is often used, where each biological element, such as a cancer cell or ECM fibre, is distinct and the interactions between the agents are defined. For example, Prasanna et al. ^26^ used a cellular Potts model-based multiscale computational framework to investigate spatial tumour heterogeneity. Finally, hybrid models combine multiple methods, such as modelling the cells using a discrete model and the substrate using a continuous model. Poonja et al. ^27^ built a model that combines an off-lattice agent-based model for the cells with a vector field representation of the ECM fibril structure. In this model, the ECM is characterised by fibril orientation and stiffness, which can inhibit cell proliferation or trigger cell migration when respective stiffness thresholds are exceeded.

Several software tools that utilise agent-based models for studying collective cell migration problems have gained popularity. Notable examples include Chaste ^28^, CompuCell3D, which uses a cellular Potts model ^29^, and PhysiCell, which employs an off-lattice centre-based agent-based model ^30^. PhysiCell has a growing user community and has been previously used to model the extracellular matrix. Gonçalves and Garcia-Aznar ^15^ developed a model for spheroid growth with PhyiCell by defining the ECM as part of the chemical microenvironment with zero diffusivity. The extracellular matrix in PhysiCell has also been modelled as an agent in a PhysiCell addon, named PhysiMeSS ^31^. Here each ECM fibre is represented by cylinders with varying stiffness. Another ECM extension of PhysiCell was developed by Metzcar et al. ^32^. They modelled the ECM as a continuum, then discretised into smaller volumetric elements which store information about the ECM fibre orientation, average anisotropy and fibre density.

We present a hybrid discrete-continuous model built upon the ECM framework developed by Metzcar et al. ^32^ in PhysiCell, utilising its ECM fibre density feature. The model explores the influence of cell-cell and cell-matrix interactions on cancer spheroid growth at different levels of ribose-induced ECM stiffness. Our model accounts for cell-cell and cell-ECM adhesion and repulsion, ECM remodelling, and cell proliferation with associated inhibition of proliferation function, allowing us to successfully replicate the experimental finding of Jahin et al. ^12^, which investigated cancer spheroid growth and invasion of non-invasive MCF7 and invasive HCC1954 cells at different ribose-induced stiffnesses of the ECM. Consistent with the *in vitro* experiments, our results indicate that ribose-induced stiffening can significantly reduce ECM remodelling and confine cancer cell movement, inhibiting spheroid growth and invasion. Moreover, this flexible modelling framework is able to incorporate additional ECM characteristics and microenvironmental conditions, such as fibre orientation and nutrient diffusion, to further refine the dynamics of cancer spheroid-ECM interaction in the future.

## 2 Materials and methods

### 2.1 Experimental data

In this paper, we aim to investigate the mechanisms involved in cancer spheroid growth and invasion into the extracellular matrix. Spheroid growth refers to the expansion of the central spheroid mass and its volume change over time as a result of cell proliferation. On the other hand, invasion describes the penetration of single cells or broad multicellular protrusions into the surrounding ECM. We build the model based on the *in vitro* experiments conducted by Jahin et al. ^12^ studying the effect of ribose-induced ECM stiffening on cancer spheroid growth and invasion of non-invasive MCF7 and invasive HCC1954 breast cancer cell lines.

In their study, Jahin et al. ^12^ formed tumour spheroids of 200 *µ*m in diameter using the hanging drop method and subsequently embedded them in a collagen matrix with varying ribose concentrations of 0 mM, 50 mM and 200 mM as in Phillips et al. ^33^. They used the non-invasive parental MCF7 and invasive parental HCC1954 cells, both human breast carcinoma cell lines with epithelial-like morphology. To model the ECM, they chose collagen I, derived from rat tail tendons, as it is the most abundant protein component in the extracellular matrix surrounding solid tumours, with supplementary fibronectin also included to allow enhanced cell attachment. During collagen hydrogel formation, ribose, a cross-linker used for non-enzymatic glycation to induce gel stiffening in *in vitro* models, was added at appropriate concentrations to increase hydrogel stiffness. More cross-linking between collagen fibres increases the ECM stiffness without altering the matrix organisation and the ligand binding sites for cell-ECM adhesion. This allows for the investigation of the effect of stiffness alone on cancer spheroid growth and invasion. The spheroids were imaged by combining Z-slices at 10 *µ*m intervals, covering the whole spheroid thickness. The images were captured daily over at least 96 hours and were used to track the spheroid invasion.

### 2.2 Model

#### 2.2.1 PhysiCell and general framework

PhysiCell is an open-source cross-platform compatible multiscale modelling tool, based in C++^30^. It employs a hybrid discrete-continuum approach, coupling an agent-based model for the cells with a continuum model for the diffusive microenvironment. The agent-based model is off-lattice and centre-based. Each agent, corresponding to a single cell, is modelled as a sphere, with its position defined by its centre. The continuum microenvironment consists of chemical substrates with associated diffusion coefficients, decay rates, sources and sinks, and initial and boundary conditions. PhysiCell is coupled to an efficient multi-substrate diffusion solver called BioFVM ^34^ to simulate the chemical microenvironment using reaction-diffusion PDEs. PhysiCell uses multiscale modelling, as it has been developed with the aim of modelling problems in cancer biology and tissue engineering, which involve processes occurring at different time scales. The system is updated using pre-defined and user-defined parameters and functions, making this tool flexible and customisable.

We build our model upon the PhysiCell (version 1.12.0) ECM framework developed by Metzcar et al. ^32^. The continuum ECM is defined separately from the chemical microenvironment and is discretised into volumetric elements, or voxels. In our model, each voxel stores information about the local ECM density. The model consists of three parts:

- ECM remodelling, corresponding to changes in ECM density due to degradation by the cells (Section 2.2.2);
- Cell movement, as a result of cell-cell and cell-ECM interactions (Section 2.2.3);
- Cell proliferation, with associated inhibition of proliferation function (Section 2.2.4).

We assume that the ribose concentration affects how the cells interact with the ECM, reducing ECM remodelling and slowing cell migration. For ECM remodelling and cell movement processes, we update the system every mechanics time step Δ*t*_*mech*_ at the default value of 0.1 min, whilst slower cell phenotype processes, *i*.*e*. cell proliferation and volume changes, are updated at a slower rate, every phenotype time step Δ*t*_*cell*_ at the default value of 6 min ^30^.

We present a constrained 3D model representing a *z*-slice image of the experimental data (Figure 1(A)). We use a single layer of ECM voxels inside which the cell agents can move freely, though their movement is restricted in the *z*-direction. The cells are spherical agents with a maximum volume *V* and corresponding radius *R* that can interact with the ECM voxels.

**Figure 1:**
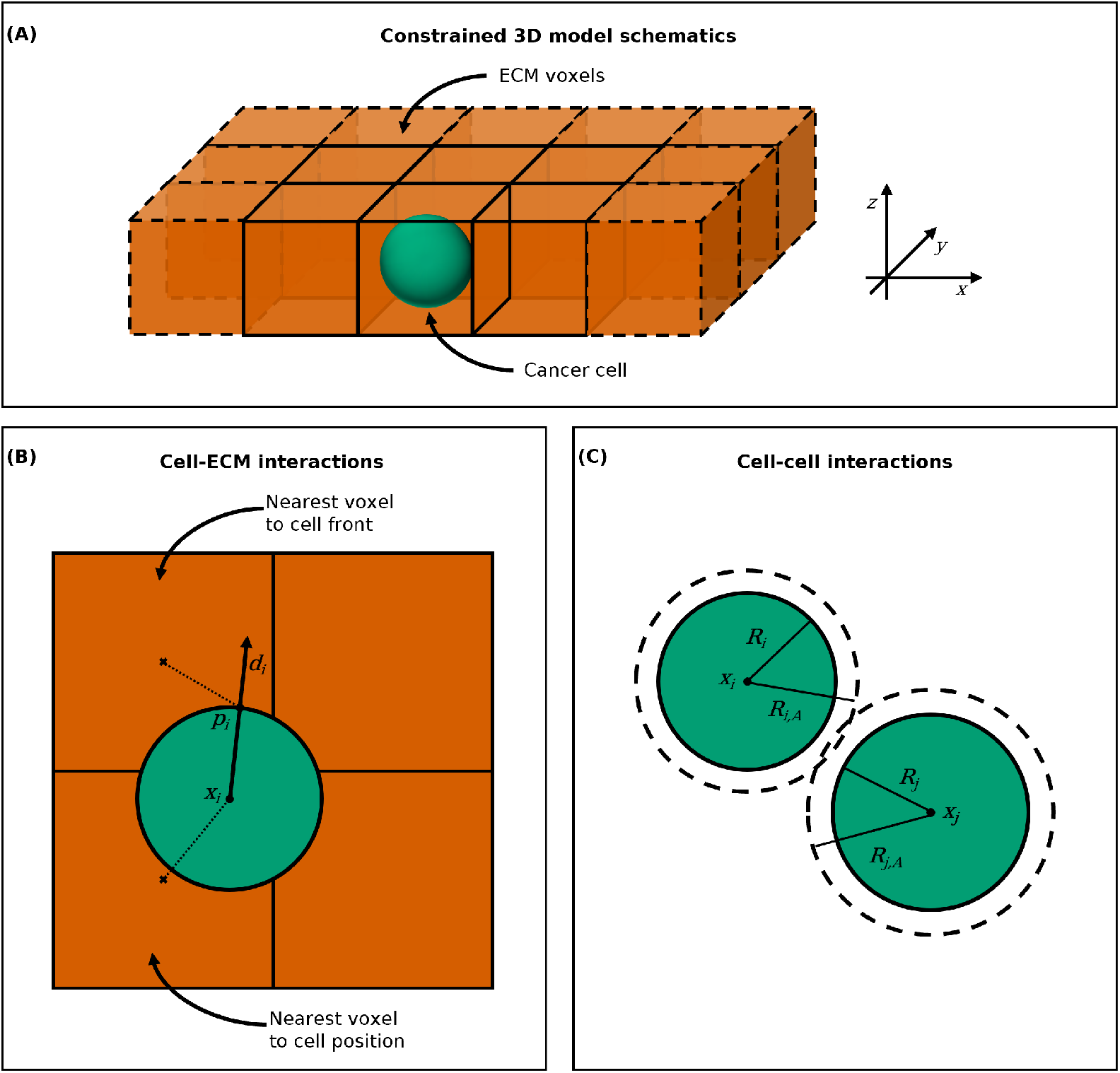
**(A)** Constrained 3D model schematics. The ECM voxels are represented as orange cubes and the cancer cell as a green sphere. The cancer cell agent moves freely within the ECM voxels in the *x*− and *y*-directions, but not in the *z*-direction. **(B)** Cell-ECM interactions computation. The cell (green) interacts with the ECM voxel (orange) whose centre (***×***) is nearest to either its position ***x***_***i***_ or its front ***p***_*i*_, determined as the point on the cell surface in its direction of movement ***d***_*i*_ **(C)** Cell-cell interactions computation. Cell *C*_*i*_ on the left has a radius *R*_*i*_ and an interaction radius *R*_*i,A*_ and cell *C*_*j*_ on the right has a radius *R*_*j*_ and an interaction radius *R*_*j,A*_. The two cells interact when their distance is less than *R*_*i,A*_ + *R*_*j,A*_.

To compute cell-ECM interactions for cell movement and ECM remodelling, we must choose which ECM voxel each cell agent interacts with. In PhysiCell the default voxel accessed for cell-substrate interactions is the nearest voxel (centre) to the position ***x***_*i*_ of the cell *C*_*i*_ (Figure 1(B)). However, invading cancer cells can form outward protrusions, such as filopodia and invadopodia, to adhere and remodel the ECM fibres mechanically and chemically ^35^. During chemical remodelling, cancer cells can use these protrusions to secrete soluble or membrane-bound matrix metalloproteinases (MMPs), a class of matrix-degrading enzymes crucial for invasion ^36^. For this reason, we introduce an alternative method of selecting the nearest voxel to the cell. When remodelling the ECM, we modify the density of the nearest ECM element to the cell front, corresponding to the nearest voxel to the point ***p***_*i*_ on the cell surface in its direction of movement ***d***_*i*_, as shown in Figure 1(B). We also use the nearest voxel to the cell front to find the local ECM density when computing the cell-ECM adhesion speed (Equation (5)). If the cell is not moving, and so ***d***_*i*_ = **0**, we select the nearest voxel to the cell position ***x***_*i*_ (Figure 1(B)).

For interactions between cell agents, such as cell-cell adhesion, as in PhysiCell, we consider the set of neighbouring cells *N*_*i*_ defined as all the cells within interaction distance *R*_*i,A*_ + *R*_*j,A*_ from the cell *C*_*i*_’s centre ***x***_***i***_ (Figure 1(C)). *R*_*i,A*_ and *R*_*j,A*_ are the maximum interaction (or adhesion) radii of the cells *C*_*i*_ and *C*_*j*_ respectively and they are fixed multiples of the cells radii ^30^.

#### 2.2.2 ECM remodelling

When a cell enters into contact with an ECM element, it remodels the matrix substrate by changing its density *ρ* ∈ [0,1]. We assume the cells degrade the ECM by dynamically reducing its density towards zero. The ECM density update equation is the following:

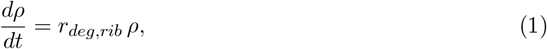

where *r*_*deg,rib*_ is the cell’s characteristic rate of degradation of the ECM, which depends on the ribose concentration. It has been observed that an increase in ribose and fibre cross-linking through glycation correlates to less ECM remodelling and degradation of the ECM fibres ^37,38^. Therefore, we assume that the ECM degradation rate *r*_*deg,rib*_ depends on the ribose concentration *rib* as follows:

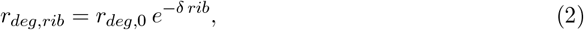

where *δ* ≥ 0 is a parameter that determines how strongly the ribose affects the cell’s base degradation rate *r*_*deg*,0_, *i*.*e*. when the ribose concentration is zero. By choosing *δ* = 0 we assume that the ribose concentration does not affect the cell’s degradation rate. If *δ >* 0, as the ribose concentration *rib* increases, the degradation rate *r*_*deg,rib*_ decreases, tending to zero as *rib* goes to infinity.

#### 2.2.3 Cell movement

The total velocity ***v***_*i*_ of a cell *C*_*i*_ can be written as

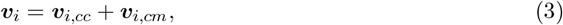

where ***v***_*i,cc*_ is the cell-cell interaction velocity as a result of cell-cell adhesion and repulsion and ***v***_*i,cm*_ is the cell-matrix interaction velocity as a result of cell-ECM adhesion and repulsion.

##### Cell-cell interactions

To reproduce cell-cell interactions, we use the built-in functions in PhysiCell for cell-cell adhesion and repulsion, in PhysiCell referred to as cell mechanics ^30,32,39^. The cell-cell interaction velocity ***v***_*i,cc*_ (Equation (3)) is a result of cell-cell adhesion and repulsion forces. When the distance between two cell centres |***x***_*j*_ − ***x***_*i*_| is less than their interaction distance *R*_*i,A*_ + *R*_*j,A*_, cell-cell adhesion is activated and the cell agents start pulling each other (Figure 1).

On the other hand, the cell-cell repulsion force is activated when two cells start overlapping, so when the distance between the two cell centres |***x***_*j*_ − ***x***_*i*_| is less than the sum of their radii *R*_*i*_ + *R*_*j*_ (Figure 1). This force is used to reproduce the effect of volume exclusion, and resistance to cell deformation when a cell is pushed by other cells.

##### Cell-matrix interactions

The cell-matrix interaction velocity ***v***_*i,cm*_ (Equation (3)) is a result of cell-ECM adhesion and repulsion

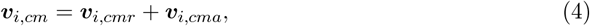

where ***v***_*i,cmr*_ is the cell-ECM repulsion velocity and ***v***_*i,cma*_ is the cell-ECM adhesion velocity.

The cell-ECM adhesion velocity ***v***_*i,cma*_ is a result of cell adhesion to the ECM fibres and can be written as

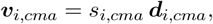

where *s*_*i,cma*_ is the cell-ECM adhesion speed and ***d***_*i,cma*_ is the cell-ECM adhesion direction. The cell-ECM adhesion direction ***d***_*i,cma*_ is given by a uniform random unit vector, whilst the cell-ECM adhesion speed *s*_*i,cma*_ depends on the ECM density *ρ*. A higher density of the ECM corresponds to a higher number of cell-ECM adhesion sites. Therefore, we define the cell-ECM adhesion speed as a linearly increasing function for cell-ECM adhesion speed with respect to the ECM density *ρ*:

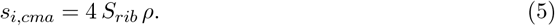

*S*_*rib*_ is the maximum cell-ECM interaction speed the cell can reach for a certain concentration of ribose *rib*. A higher ribose concentration corresponds to more cross-linking between the collagen fibres. This affects the mechanical remodelling of the fibres, as it makes it harder for the cells to realign the ECM fibres, which is essential to allow direct cell migration and invasion ^40^. Since we do not have fibre alignment and orientation in our model, we assume that higher ribose concentration, and collagen stiffness, relate to slower cell migration. Thus, we further assume that *S*_*rib*_ decreases as the ribose concentration increases as follows

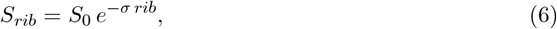

where *σ* ≥ 0 is a parameter that determines how strongly the ribose affects the maximum cell-ECM interaction speed at ribose concentration 0 mM (*S*_0_). Similarly to the role of *δ* in Equation (2), by setting *σ* = 0 we assume that the ribose concentration does not affect the cell maximum speed, while if *σ >* 0 the cell maximum speed decreases as the ribose concentration *rib* increases, tending to zero as *rib* goes to infinity.

Further, in a 3D matrix, the ECM fibres act as an obstacle to cell migration when the matrix is dense. When the ECM density *ρ* is equal to 1 the cells will be fully repelled by the ECM, which will act as a wall, and when *ρ* is equal to zero there is no repulsion. Therefore, we define the ECM density-dependent cell-ECM repulsion velocity as

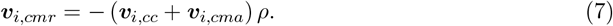

Equations (5) and (7) indicate that as the ECM density increases, both cell-ECM adhesion and repulsion speeds increase, leading to a non-monotonic resultant total cell-matrix speed (*s*_*i,cm*_). Assuming ***v***_*i,cc*_ = **0**,

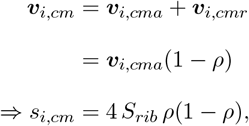

which reaches its maximum at *ρ* = 0.5, where *s*_*i*_ = *S*_*rib*_.

##### Persistence time

In PhysiCell ^30^, the cell-ECM adhesion velocity ***v***_*cma*_ changes stochastically every mechanical time step Δ*t*_*mech*_ with probability

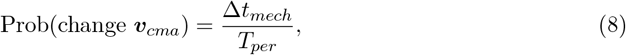

where *T*_*per*_ is the persistence time. However, the persistence in cell movement is defined as the mean time a cell maintains its direction of motion, not its speed ^41^. Therefore, we update the cell-ECM adhesion direction ***d***_*i,cma*_ and the cell-ECM interaction speed *s*_*i,cma*_ separately. The cell-ECM adhesion direction changes stochastically with the same probability as in Equation (8), whilst the cell-ECM interaction speed gets updated deterministically every mechanics time step. In this way, the cell is able to rapidly react to changes in the ECM density and tune its speed accordingly.

#### 2.2.4 Cell proliferation

For cell proliferation, we use a live cell model from PhysiCell ^30^. This simple model for proliferation consists of cells dividing in any time interval [*t, t* + Δ*t*] with probability:

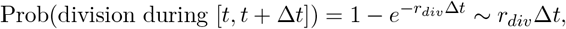

where *r*_*div*_ is the cell proliferation (or division) rate. When dividing, the cell will halve its volume, duplicate the cell with all its state and parameter values and place the daughter cells side by side with their centres inside the radius of the original cell. The daughter cells then grow in volume until reaching the maximum volume *V*.

However, compression of the tumour spheroid due to confinement and lack of nutrients can slow or arrest cell proliferation ^42–44^. The spheroid can be compressed when the surrounding ECM is too dense and is not degraded quickly enough, slowing proliferation ^42^. Nutrient diffusion depends on the porosity of the ECM, which in turn depends on the density of the fibres ^44^. Furthermore, the cells in the centre of the spheroid are less exposed to nutrients, since the cells in the outer layer consume the nutrients first ^45^. Therefore, as we chose not to include nutrient diffusion in the current model, we simplify inhibition of proliferation by assuming that proliferation is inhibited when the cells are surrounded by neighbours (the number of neighbours above a pre-defined overcrowding threshold) and is slowed down by the presence of extracellular matrix. We rewrite the probability of division in any time interval [*t, t* + Δ*t*] as

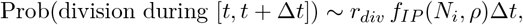

where *f*_*IP*_ is the inhibition of proliferation function defined as

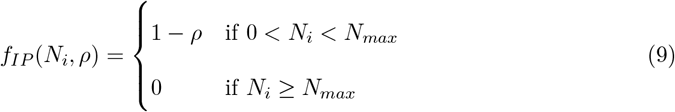

with *N*_*i*_ being the number of neighbours of the cell *C*_*i*_, *N*_*max*_ the overcrowding threshold and *ρ* ∈ [0,1] the ECM density of the nearest voxel to the cell position.

### 2.3 Statistical analysis

To compare our results with the experimental data, we calculated spheroid area growth relative to the initial time, cell count and Delaunay mean distance between cells in Python version

3.10.12. Given the stochastic nature of our model, we ran 10 replicates for each simulation and computed the mean and 25th/75th percentile of spheroid area growth relative to the initial time, cell count and Delaunay mean distance every 60 min.

We computed the spheroid area growth relative to the initial time by calculating the area covered by the cells at each time point and dividing it by the area covered at the initial time *t* = 0 min. The spheroid area was approximated by dividing the entire domain into a 5000*×*5000, initially setting all grid elements to a value of 0. This baseline value represents unoccupied space. We then drew disks of value 1 at the coordinates of each cell’s centre with their corresponding radius (Figure 2(A)). Overlaps were ignored, as grid elements covered by multiple cells are only counted once. To calculate the total spheroid area, we summed the grid elements with value 1 and rescaled to the original domain size to obtain the spheroid area in *µ*m^2^. This process is analogous to the method used for computing spheroid invasion relative to the initial time in Jahin et al. ^12^.

**Figure 2:**
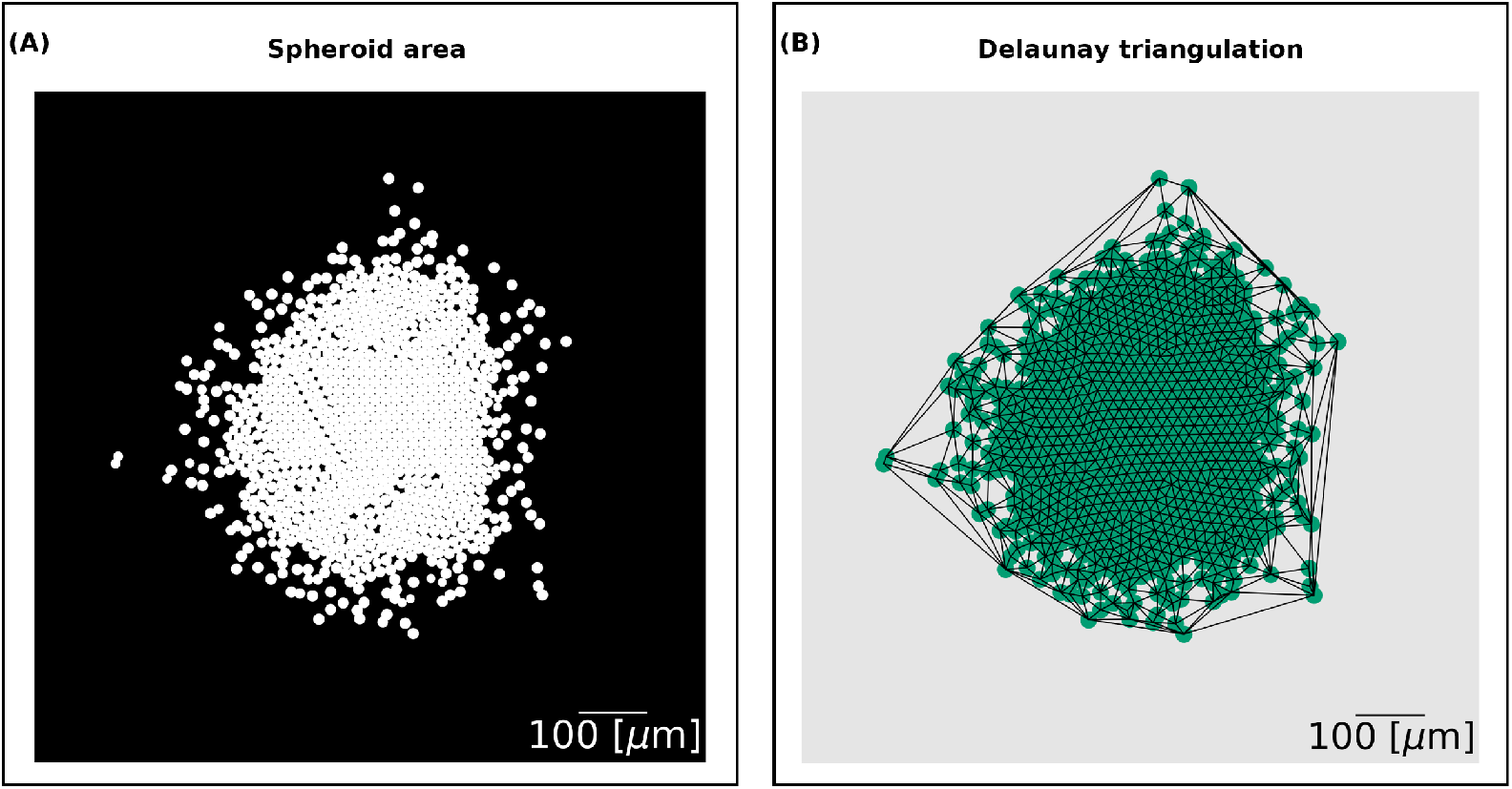
**(A)** Spheroid area output. The domain is divided into a 5000*×*5000 grid. The grid elements that overlap with a cell have value 1 and are shown in white. The grid elements corresponding to the background have value 0 and are shown in black. The spheroid area is computed by summing together the elements of the grid and rescaling to the original size of the domain. **(B)** Delaunay triangulation output. We generate a network that connects the centres of the cells using the spacial algorithm Delaunay from the Python library SciPy. The scale bar on the bottom right is of length 100 *µ*m.

For the cell count, we tallied the total number of cells in the simulations at each time point, corresponding to the number of nuclei in a slice of the experimental data.

Finally, the Delaunay mean distance measures the proximity of cells. It utilises the Delaunay triangulation, the dual graph of the Voronoi diagram, of a set of points. The edges of the graph form triangles whose circumscribed circles do not contain any other point. We used the cell centres as the input nodes of the network and generated the Delaunay network using the spacial algorithm Delaunay from the Python library SciPy version 1.11.1 (Figure 2(B)). We then computed the mean edge length between the nodes to find the Delaunay mean distance.

## 3 Results

### 3.1 Impact of key cell-ECM interaction parameters on spheroid growth

Invasion and migration of cancer cells from a cancer spheroid into the surrounding extracellular matrix depends strongly on how they interact with the ECM ^3^. The cancer cells need to remodel the ECM to enable invasion, and can then use the adhesion sites on the collagen fibres to propel themselves and invade further. In our model, to study such cell-ECM interactions, we consider different biophysical parameters representing cell-ECM cross-talk and specific properties of the ECM. We control how quickly the agent cells reduce locally the ECM density by changing the degradation rate (*r*_*deg*,0_, Equation (1)). In turn, the ECM density affects the cell’s speed (Equations (5) and (7)), which reaches its maximum cell-ECM interaction speed (*S*_*rib*_) when the ECM density is equal to 0.5. The ECM density also affects the cell’s proliferation rate (*r*_*div*_) through the inhibition of proliferation function (Equation (9)). In this section, we present the results of our analysis on the degradation rate (*r*_*deg*,0_) and maximum cell-ECM interaction speed (*S*_0_) without any ribose at different proliferation rates (*r*_*div*_). Then we analyse the *δ* and *σ* parameters, which determine how strongly ribose affects the degradation rate (*r*_*deg,rib*_, Equation (2)) and the maximum cell-ECM interaction speed (*S*_*rib*_, Equation (6)) respectively at different ribose concentrations (*rib*).

We initiated all simulations with a spheroid of cancer cells of 200 *µ*m in diameter and homogeneous ECM density with value *ρ* = 1 throughout, except at the spheroid’s location, where the ECM density is zero. We set the overcrowding threshold *N*_*i*_ used in the inhibition of proliferation function (Equation (9)) to 6, equivalent to a cell fully surrounded by the other cells. All other parameters are set to their default values as used in PhysiCell 1.12.0^30^.

We examined the impact of the degradation rate (*r*_*deg*,0_) and the maximum cell-ECM interaction speed (*S*_0_) on spheroid area growth relative to the initial time and Delaunay mean distance (explained in Section 2.3), holding the ribose concentration at 0 mM. The analysis was conducted for three proliferation rates: *r*_*div*_ = 0.0004 min^−1^, 0.0006 min^−1^ and 0.0008 min^−1^ (Figure 3(A), (B)).

**Figure 3:**
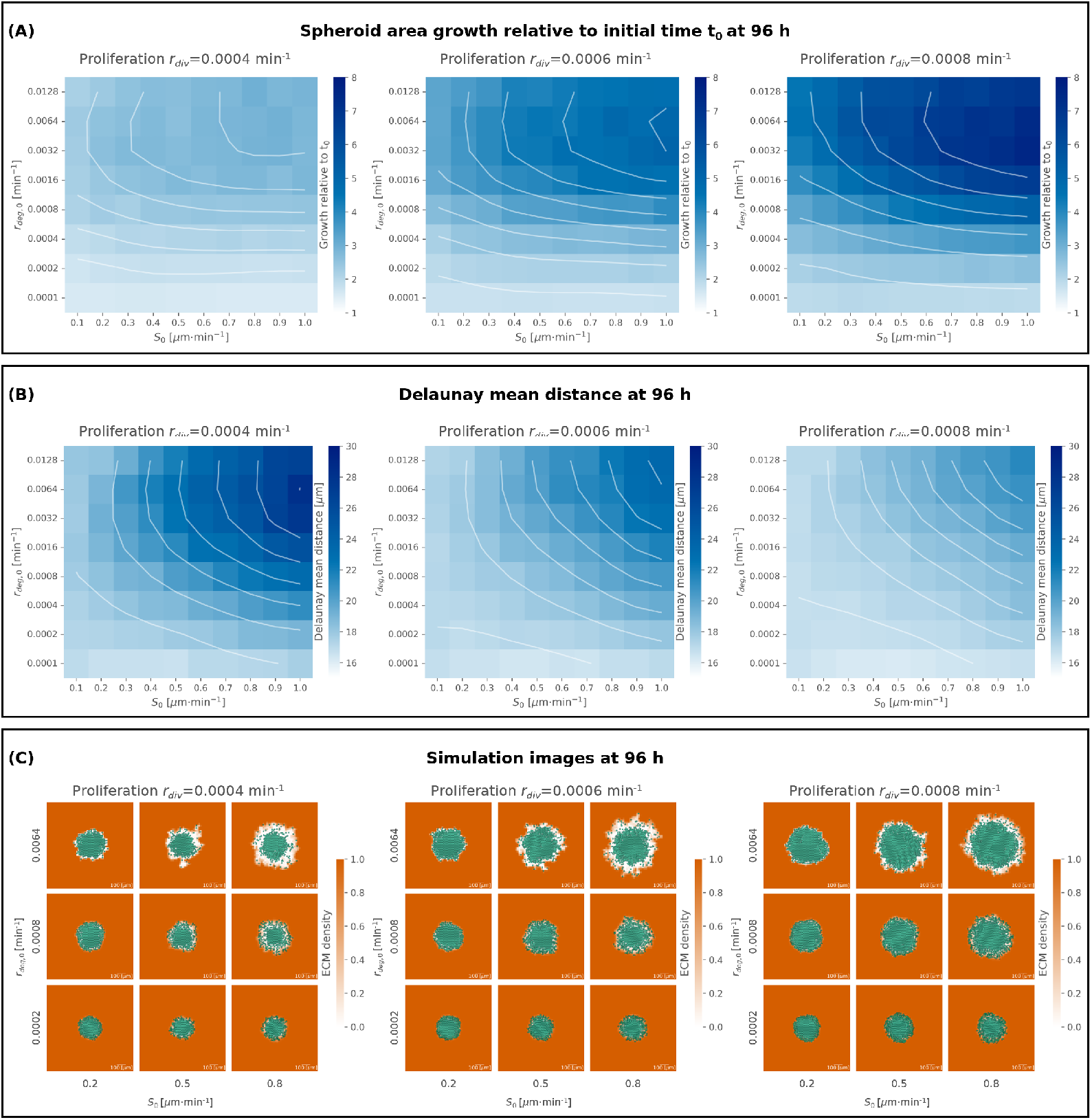
**(A, B)** Heatmaps showing the effects on spheroid area growth relative to the initial time *t*_0_ **(A)** and Delaunay mean distance **(B)** after 96 h with ribose concentration 0 mM of proliferation rate *r*_*div*_ (columns), maximum cell-ECM interaction speed *S*_0_ (*x*-axis) and degradation rate *r*_*deg*,0_ (*y*-axis). The colour intensity represents the mean values over 10 replicates of the spheroid area growth relative to the initial time ranging from 1 to 8 and Delaunay mean distance ranging from 10 *µ*m to 30 *µ*m, as shown in the colour bars. Contour lines are also shown in white. **(C)** Tables showing the simulation figures at 96 h for varying proliferation rate *r*_*div*_ (columns), maximum cell-ECM interaction speed *S*_0_ (*x*-axis) and degradation rate *r*_*deg*,0_ (*y*-axis). The cells are represented in semi-transparent green and the ECM density is in orange taking values between 0 and 1, as shown in the colour bar.

Our results demonstrate that an increase in proliferation rate (*r*_*div*_) enhances spheroid area growth relative to the initial time (Figure 3(A)) and reduces the Delaunay mean distance (Figure 3(B)). This is expected, as faster cell division leads to a denser cell population, which is correlated to a lower average cell-cell distance.

Further, increasing the maximum cell-ECM interaction speed (*S*_0_) leads to enhanced spheroid growth and a higher Delaunay mean distance (Figure 3(A), (B)). A higher *S*_0_ allows more cells to detach from the spheroid, reducing the number of neighbours, and consequently avoiding proliferation arrest. It also enables cells to access and degrade more areas of the ECM, further promoting proliferation. Additionally, with faster cell migration, cells at the spheroid’s edge become more dispersed, contributing to the increase in Delaunay mean distance.

Finally, we observed that increasing the degradation rate *r*_*deg*,0_ does not always induce a monotonic increase in spheroid area growth relative to the initial time and Delaunay mean distance. Generally, increasing the degradation rate enhances spheroid growth (Figure 3(A)) and increases cell-cell distance, leading to a higher Delaunay mean distance (Figure 3(B)). This occurs because a higher degradation rate reduces ECM density around the spheroid, which in turn positively affects proliferation and invasion. However, when the degradation rate is excessively high (*r*_*deg*,0_ = 0.0128 min^−1^), the ECM is degraded too quickly, which inhibits migration and invasion. Therefore, for fixed proliferation rate *r*_*div*_ and maximum cell-ECM interaction speed *S*_0_, as the degradation rate *r*_*deg*,0_ increases, the spheroid growth slows down (Figure 3(A)) and the Delaunay mean distance decreases (Figure 3(B)).

Figure 3(C) shows simulation images at the final time point (96 h) for *S*_0_ = 0.2, 0.5 and 0.8 *µ*m*·*min^−1^, *r*_*deg*,0_ = 0.0002, 0.0008 and 0.00064 min^−1^ and *r*_*div*_ = 0.0004, 0.0006 and 0.0008 min^−1^. These images illustrate that higher *S*_0_ values lead to greater cell dispersion at the spheroid’s edge. When combined with higher ECM degradation *r*_*deg*,0_ more single cells are observed migrating away from the spheroid. Thus, increased degradation rate and maximum cell-ECM interaction speed contribute to the formation of protrusions in the spheroid. In contrast, lower degradation levels and migration speed limit spheroid growth, resulting in a more rounded spheroid shape.

We then analysed the impact of the parameters *δ* and *σ* on spheroid area growth relative to the initial time (Figure 4). For *δ, σ* = 0 the functions are constant, so *r*_*deg,rib*_ = *r*_*deg*,0_ and *S*_*rib*_ = *S*_0_, while for *δ, σ >* 0 the functions are monotonically decreasing with respect to the ribose concentration *rib*, so *r*_*deg,rib*_ *< r*_*deg*,0_ and *S*_*rib*_ *< S*_0_ for ribose *rib* greater than zero (Equations (2) and (6)). For this analysis, we fixed the degradation rate at ribose concentration 0 mM (*r*_*deg*,0_ = 0.0032 min^−1^), the maximum cell-ECM interaction speed at ribose concentration 0 mM (*S*_0_ = 0.7 *µ*m*·*min^−1^) and the proliferation rate (*r*_*div*_ = 0.00072 min^−1^). The analysis was conducted for two ribose concentrations: *rib* = 50 mM and 200 mM.

**Figure 4:**
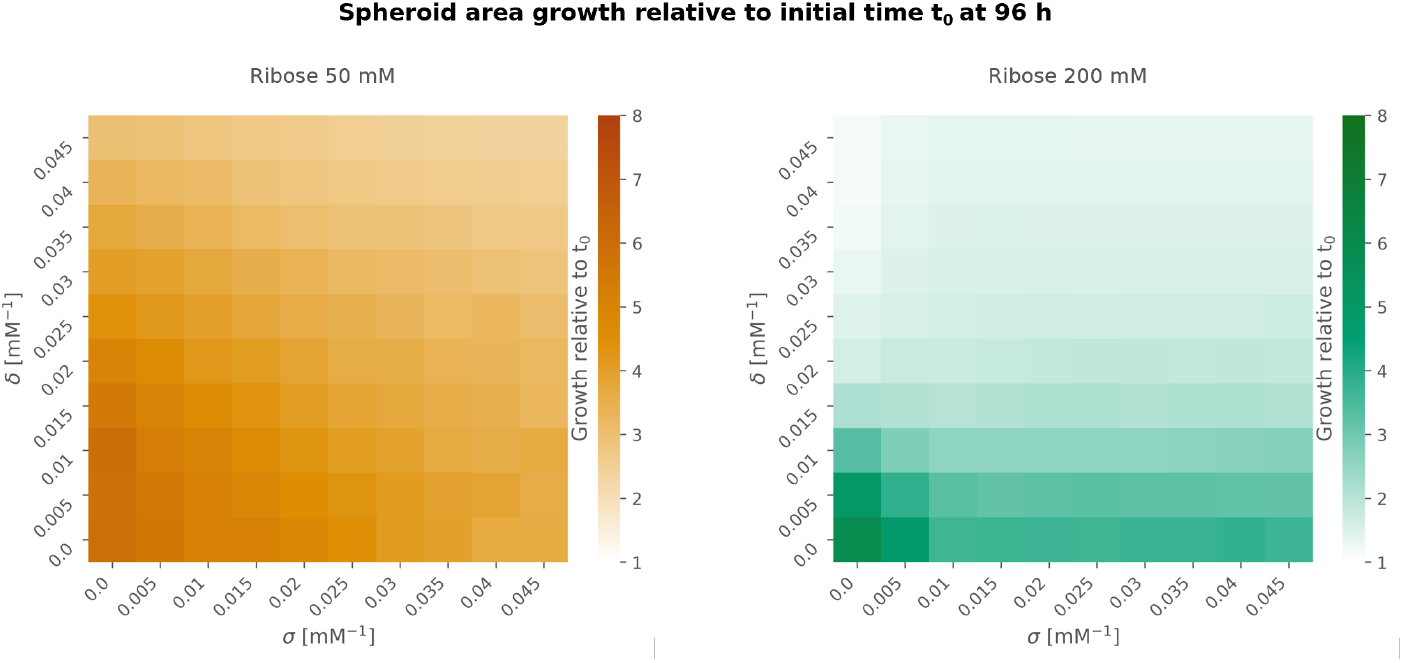
Heatmaps showing spheroid growth relative to the initial time *t*_0_ for ribose concentration of 50 mM in orange (left) and 200 mM in green (right) for varying values of the parameters *δ* (*y*-axis) and *σ* (*x*-axis). We set *r*_*deg*,0_ = 0.0032 min^−1^, *S*_0_ = 0.7 *µ*m*·*min^−1^ and *r*_*div*_ = 0.00072 min^−1^. The colour intensity represents the mean values over 10 replicates of the spheroid area growth relative to the initial time ranging from 1 to 8, as shown in the colour bars.

Figure 4 shows that higher ribose concentrations correspond to a decrease in spheroid growth, and increasing either *δ* or *σ* also results in reduced spheroid growth. Notably, at 200 mM ribose, when *σ* is greater than or equal to 0.015 mM^−1^, spheroid growth remains unchanged for fixed values of *δ*. This occurs because for *σ* = 0.015 mM^−1^ the maximum cell-ECM interaction speed is significantly reduced, *S*_200_ = *S*_0_ *e*^−*σ* 200^ ∼ 0.035 *µ*m*·*min^−1^ compared to *S*_50_ = *S*_0_ *e*^−*σ* 50^ ∼ 0.33 µm·min^−1^. As a result, as the cell population becomes denser, proliferation is inhibited throughout the spheroid except at its boundary, where proliferation depends on the surrounding ECM density. Consequently, spheroid growth is determined by the rate at which cells degrade the ECM at the boundary, facilitating increased proliferation in this region.

### 3.2 Model captures inhibition of cancer spheroid growth of non-invasive and invasive breast cancer cells when increasing ribose concentration

To begin with, we replicated experiments from Jahin et al. ^12^ that study the effect of ribose concentration on two different cell lines of parental breast cancer cells: MCF7 and HCC1954. MCF7 cells are a non-invasive cell line, which correlates with weaker cell-ECM interactions ^46^. On the other hand, HCC1954 cells are a more aggressive and invasive cell line, characterised by enhanced contractility, and therefore stronger interactions with the ECM fibres enabling migration, and further ECM remodelling ^12,47^. The *in vitro* experiments showed that with increasing ribose concentration, and therefore collagen fibre stiffness, the spheroid invasion was inhibited for the invasive HCC1954 cells, while the non-invasive MCF7 cells did not invade for any of the ribose concentrations.

The cell-ECM interactions in our model depend on two parameters: ECM degradation rate, which controls the ECM remodelling by the cells (Equation (1)), and maximum cell-ECM interaction speed, which affects the cell’s movement (Equation (5)). Given the different invasiveness of the two cell lines, we assume that the invasive cells have a higher ECM degradation rate and maximum cell-ECM interaction speed than the non-invasive cells. From the wide range of values studied in Section 3.1 (Supplementary Figure S1), we choose the values that lead to simulations matching the experimental observations (Figure 5(E), (F)). Hence, we use the following set of values: ECM degradation rate *r*_*deg*,0_ = 0.0001 min^−1^ and maximum cell-ECM interaction speed *S*_0_ = 0.1 *µ*m*·*min^−1^ for non-invasive cells, and ECM degradation rate *r*_*deg*,0_ = 0.0032 min^−1^ and maximum cell-ECM interaction speed *S*_0_ = 0.7 *µ*m*·*min^−1^ for invasive cells. Further, the strength of the effect of ribose concentration on the cell behaviour depends on the parameters *δ* for degradation rate *r*_*deg,rib*_ (Equation (2)) and *σ* for maximum cell-ECM interaction speed *S*_*rib*_ (Equation (6)). Following Section 3.1 (Figure 4), we use *δ* = 0.02 mM^−1^ and *σ* = 0.035 mM^−1^ to match the spheroid growth after 96 h of the invasive cells for ribose concentrations of 50 mM and 200 mM (Figure 5(D)(i)). Finally, we set the proliferation rate *r*_*div*_ = 0.00072 min^−1^, which is the default parameter for the live cell cycle model of MCF10A breast cancer epithelial cells in PhysiCell. ^30^ Following the *in vitro* experiments performed in Jahin et al. ^12^, we set the initial spheroid diameter to be 200 *µ*m (corresponding to 139 cells), with homogeneous ECM density with value *ρ* = 1 throughout, except at the spheroid’s location, where the ECM density is zero. We study the effect of ribose concentrations, 0 mM, 50 mM and 200 mM, on spheroid area growth relative to the initial time, cell count and Delaunay mean distance, as shown in Figure 5(C), (D).

**Figure 5:**
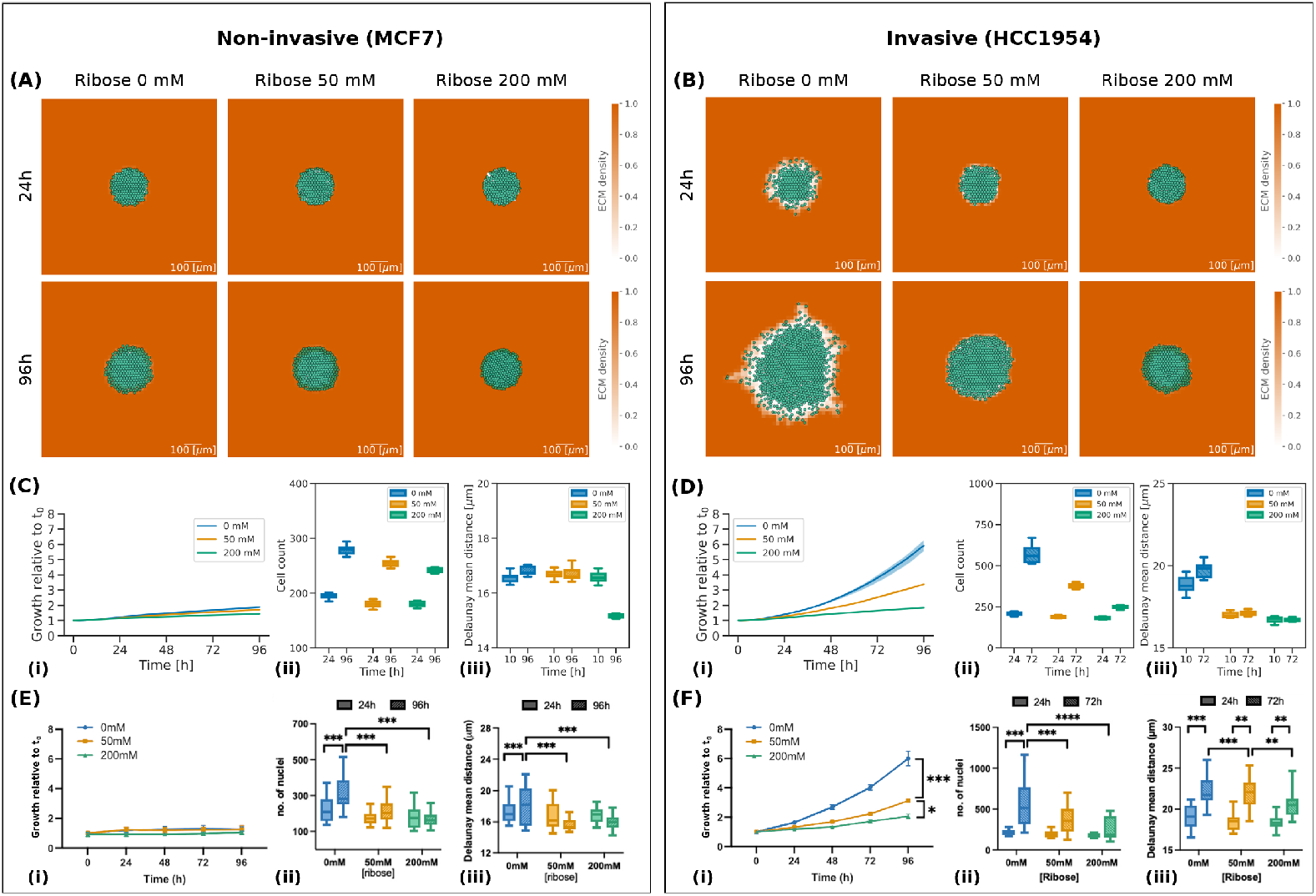
Results of simulations for non-invasive (MCF7) and invasive (HCC1954) cell lines. **(A, B)** Simulation images of non-invasive **(A)** and invasive **(B)** cell lines respectively, with ribose concentrations of 0 mM, 50 mM and 200 mM at 24 h and 96 h. The cells are represented in semi-transparent green and the ECM density is in orange, with values between 0 and 1 as indicated in the colour bar. The scale bar at the bottom right is 100 *µ*m in length. Full videos of non-invasive and invasive spheroids with ribose concentration 0 mM can be found in the Supplementary material. **(C, D)** Simulation results for non-invasive **(C)** and invasive **(D)** cell lines showing line plots of spheroid growth relative to the initial time *t*_0_ change over time **(i)**, box plots of cell count **(ii)** and box plots of Delaunay mean distance **(iii)**. The ribose concentrations are represented in blue for 0 mM, orange for 50 mM and green for 200 mM. Mean and 25th/75th percentile are shown, with the addition of min/max in box plots. **(E, F)** Adapted with permission from Jahin et al. ^12^. Plots for MCF7 (non-invasive) **(E)** and HCC1954 (invasive) **(F)** cell lines showing growth relative to the initial time *t*_0_ (i), number of nuclei (ii) and Delaunay mean distance (iii). Mean and 25th/75th percentile are shown, with the addition of min/max in box plots. Asterisks indicate statistical significance following ANOVA testing with Sidak’s post-hoc test (** = *p <* 0.01, *** = *p <* 0.001, **** = *p <* 0.0001).

In their experiments with non-invasive MCF7 cells, Jahin et al. ^12^ observed that increasing ribose concentration did not affect spheroid growth (Figure 5(E)(i)), consistent with previous reports ^48^. Our findings also indicate that spheroid growth was inhibited for the non-invasive cells, with the spheroid area growth at the final time point remaining below 2 for all ribose concentrations (Figure 5(C)(i)). However, while the invasion of MCF7 cells was low, Jahin et al. ^12^ observed an increase in the number of nuclei, particularly at 0 mM ribose (Figure 5(E)(ii)). Similarly, our simulations showed a larger increase in cell count at 0 mM ribose, with the cell count reaching ∼ 280 after 96 h (Figure 5(C)(ii)), which matches the corresponding mean number of nuclei observed *in vitro* (Figure 5(E)(ii)). Interestingly, they found that the Delaunay mean distance increased for ribose concentration of 0 mM and it decreased at 50 mM and 200 mM, indicating a denser spheroid for higher ribose concentrations (Figure 5(E)(iii)). In our simulations, we observed that the Delaunay mean distance between 10 h and 96 h slightly increased at ribose concentration 0 mM and decreased for ribose 200 mM, but remained constant at 50 mM (Figure 5(C)(iii)). This means that, in our simulations, the spheroid at 50 mM is less dense than in the experimental data. This can be due to more rapid degradation of the ECM or slower cell proliferation than in the experiments.

In contrast, the invasive HCC1954 cells exhibited a reduction in spheroid growth and the number of nuclei with a ribose concentration increase *in vitro* (Figure 5(F)(i), (ii)). Our simulations closely matched the experimental data, resulting in a spheroid growth of ∼ 6 after 96 h and a cell count of ∼ 570 after 72 h for ribose 0 mM, spheroid growth of ∼ 3.4 after 96 h with a cell count of ∼ 370 after 72 h for ribose 50 mM, and spheroid growth of ∼ 1.8 after 96 h and a cell count of ∼ 250 after 72 h for ribose 200 mM (Figure 5(D)(i), (ii)). Further, Jahin et al. ^12^ found that the Delaunay mean distance at 72 h was higher than that at 24 h, for all ribose concentrations considered. However, it decreased with an increase in the ribose concentration (Figure 5(F)(iii)). Our simulations showed that the Delaunay mean distance increased between 10 h and 72 h for ribose concentration of 0 mM while it almost stayed constant for 50 mM and 200 mM (Figure 5(D)(iii)). However, by looking at the Delaunay mean distance over time (Supplementary Figure S2) we see that the Delaunay mean distance does not monotonically increase or decrease over time. This suggests that two time points are insufficient to interpret changes over time in the compactness of the spheroids.

With our choice of parameters, we observe that low degradation rate and maximum cell-ECM interaction speed inhibit both invasion and proliferation of the spheroid. This is mainly due to the cell-ECM repulsive velocity ***v***_*cmr*_ (Equation (7)) and the inhibition of proliferation function (Equation (9)). As observed in the parameter analysis in Section 3.1, low degradation rate and maximum speed make the spheroid denser thanks to the ECM acting as a wall because of the repulsive velocity. Proliferation is inhibited at the centre of the spheroid due to the high number of neighbours and at the edge due to the high ECM density. The chosen values for *δ* and *σ* in our simulations give low degradation rates and maximum cell-ECM interaction speeds for the invasive cells at ribose concentrations of 50 mM and 200 mM. The degradation rates at ribose concentrations 50 mM and 200 mM are *r*_*deg*,50_ ∼ 0.001 min^−1^ and *r*_*deg*,200_ ∼ 0.00006 min^−1^, while the maximum cell-ECM interaction speeds are *S*_50_ ∼ 0.1 *µ*m*·*min^−1^ and *S*_200_ ∼ 0.0006 *µ*m*·*min^−1^. This indicates that both ECM degradation and cell speed are substantially reduced for the invasive cells as the ribose concentration increases, which lowers tumour invasion and proliferation. Thus, our simulations validate the observation by Jahin et al. ^12^, that ribose-induced cross-linking of collagen possibly reduces ECM remodelling and migration, slowing spheroid growth and invasion.

### 3.3 MMPs inhibition for invasive cells inhibits spheroid area growth

Cancer cells remodel the extracellular matrix mechanically and proteolytically when invading, creating paths that facilitate the migration of nearby attached cancer cells ^49^. The cells mechanically apply forces to the ECM fibres by pushing or pulling the fibres when adhering to ligand binding sites, resulting in fibre displacement and orientation changes. Fibre orientation can direct migration, and the realignment of the collagen fibres has been associated with higher invasion ^40^. On the other hand, proteolytic remodelling involves enzymatic degradation of ECM fibres through the activity of matrix metalloproteinases (MMPs). It has been shown that ECM degradation by MMPs significantly contributes to cell invasion as it facilitates migration and realignment of the fibres ^36^. Jahin et al. ^12^ investigated the role of MMPs in fibre alignment and invasion by treating the invasive HCC1954 cells with the pan-MMP inhibitor GM6001. They observed that MMP inhibition significantly reduces invasion for ribose 0 mM, but not for ribose 50 mM and 200 mM (Figure 6(C)).

**Figure 6:**
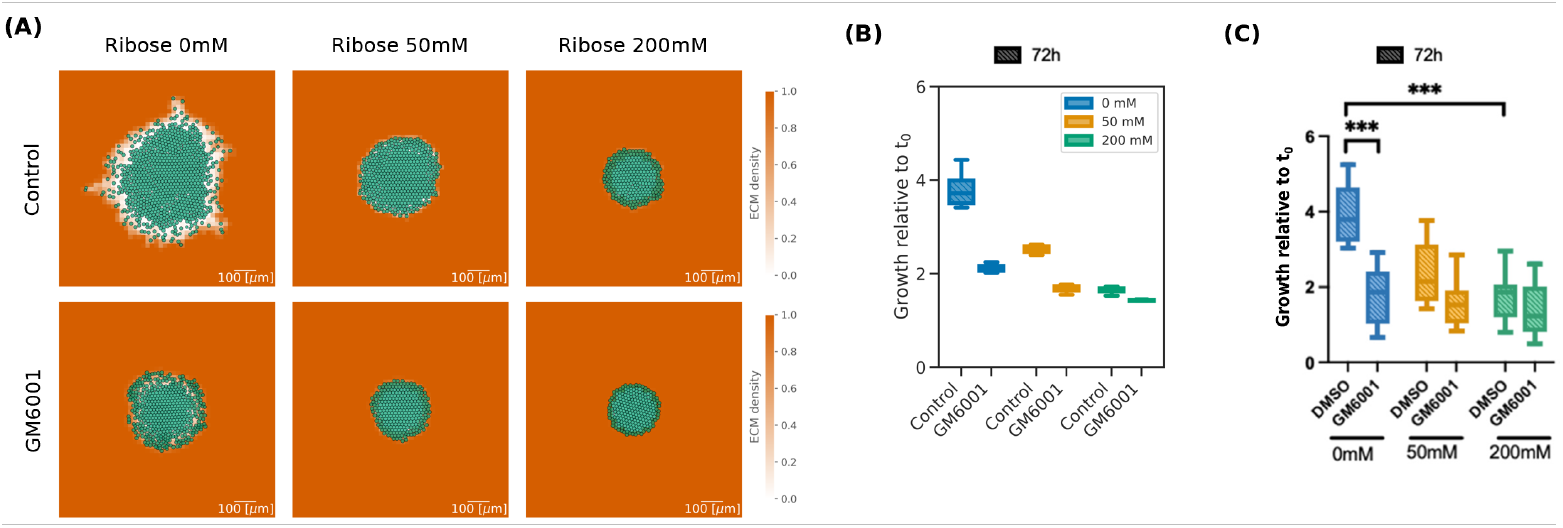
Simulations for invasive (HCC1954) cells with MMP inhibitor GM6001. **(A)** Simulation images of invasive cells with high (Control) and low (GM6001) degradation rates, with ribose concentrations of 0 mM, 50 mM and 200 mM, at 96 h. The cells are represented in semi-transparent green and the ECM density is in orange taking values between 0 and 1 as shown in the colour bar. The scale bar on the bottom right of length 100 *µ*m. **(B)** Box plots of spheroid growth relative to the initial time *t*_0_ of invasive cells with high (Control) and low (GM6001) degradation rates. The ribose concentrations are represented in blue for 0 mM, orange for 50 mM and green for 200 mM. Mean and 25th/75th percentile with min/max are shown. **(C)** Adapted with permission from Jahin et al. ^12^. Box plots of growth relative to the initial time *t*_0_ of invasive cells (HCC1954) at 72 h. Comparison between control condition (DMSO) and MMP inhibitor treatment (GM6001) at ribose concentrations 0 mM (blue), 50 mM (orange) and 200 mM (green). Mean and 25th/75th percentile with min/max are shown. Asterisks indicate statistical significance following ANOVA testing with Sidak’s post-hoc test (*** = *p <* 0.001).

In our model, ECM remodelling is represented by the degradation rate parameter *r*_*deg*,0_, which is responsible for changes in ECM density. We compared the experimental results in control conditions (DMSO) with the invasive cells simulations discussed in Section 3.2. Following the parameter analysis from Section 3.1 (Supplementary Figure S1), we choose the degradation rate matching the experimental results of growth relative to the initial time after 72 h of the invasive HCC1954 cell line with the addition of the pan-MMP inhibitor GM6001 (Figure 6(C)). We selected the degradation rate *r*_*deg*,0_ = 0.0004 min^−1^. This non-zero value arises because the MMP inhibitor only blocks ECM degradation, but not mechanical remodelling of the fibres, which makes it possible for the cells to locally change the density of the ECM even without degrading the fibres.

Similarly to the results from the experimental data by Jahin et al. ^12^ shown in Figure 6(C), spheroid area growth relative to the initial time after 72 h was significantly reduced for ribose 0 mM going from ∼ 4 in the control conditions simulation (Control) to ∼ 2 with the reduced degradation rate (GM6001) (Figure 6(B)). At ribose concentration of 50 mM spheroid growth was less affected by the MMP inhibition, both in *in vitro* and in the simulations. Finally, at 200 mM, both the Control and the reduced degradation rate (GM6001) conditions have similar spheroid sizes in the simulations after 72 h, which is in line with the experimental data in the DMSO and GM6001 conditions (Figure 6(C)).

## 4 Discussions

In this paper, we presented a hybrid discrete-continuous model built in PhysiCell (version 1.12.0) to describe the interactions between cells modelled as discrete agents and the extracellular matrix as a continuum. Our model incorporates cell-cell and cell-ECM adhesion and repulsion, ECM remodelling, and cell proliferation with associated inhibition of proliferation function, allowing us to investigate the critical role of cell-ECM interactions in cancer spheroids.

Our findings indicate that increased cell-ECM adhesion promotes invasion, while ECM degradation significantly influences spheroid growth. Notably, we observed a non-monotonic effect of ECM degradation: increasing degradation enhances growth due to reduced matrix confinement, yet excessive degradation limits migration for the cells at the edge of the spheroid, ultimately restricting invasion. Typically cells don’t over-degrade the ECM as the relationship between adhesion and cell survival is crucial. Anoikis, a programmed cell death mechanism in anchorage-dependent cells, highlights the necessity for ECM attachments since the communication between proximal cells and between cells and ECM provide essential signals for growth or survival ^50^. However, it has been found that cancer cells undergoing EMT can acquire anoikis resistance ^50^. In our model, we assume that the target value for ECM degradation is zero density (Equation (1)). This can potentially be a limitation of our model leading to wrong predictions for high degradation rates. Finally, we found that lower rates of ECM degradation and migration speeds result in more symmetrical and compact spheroids. In contrast, higher degradation and migration speeds lead to increased cell detachment and protrusion formation at the spheroid’s edge.

We replicated the experiments carried out by Jahin et al. ^12^ that investigated the impact of ribose-induced collagen stiffening on the invasion of two parental breast cancer cell lines: the non-invasive MCF7 and the invasive HCC1954. We differentiated the cell lines based on their cell-ECM interaction parameters: ECM degradation rate (*r*_*deg,rib*_), which controls the ECM remodelling by the cells (Equation (1)), and maximum cell-ECM interaction speed (*S*_*rib*_), which affects the cell’s movement (Equation (5)). We assigned low cell-ECM interaction parameter values to the non-invasive cells and high cell-ECM interaction parameter values to the invasive cells. Assuming that higher ribose concentrations reduce ECM degradation and migration speed, our model successfully predicted a decrease in spheroid area growth with increasing ribose concentration, in line with the experimental observations of Jahin et al. ^12^. Furthermore, we confirmed that inhibiting ECM degradation reduced spheroid growth in the invasive cell line.

In our current model, we represent the collagen fibre matrix as a homogeneous density and treat the ribose as a separate quantity that directly impacts ECM remodelling and cell migration. However, a more comprehensive framework of the matrix would benefit from incorporating additional ECM properties, such as fibre orientation, alignment and cross-linking. ^31,32^ Fibre orientation and alignment affect the directed migration of cells, which is a process correlated with enhanced spheroid invasion. Furthermore, as cells dynamically remodel the ECM, fibre orientation and alignment change not only locally but also at greater distances. The addition of ribose increases cross-linking between fibres and impacts both the chemical and mechanical remodelling of the ECM by cancer cells. Furthermore, cancer cells also contribute to ECM deposition and cross-linking. Our modelling framework is adaptable and allows for the integration of these additional ECM properties, such as fibre orientation and alignment, as in Metzcar et al. ^32^. In subsequent phases of the model development, we plan to implement these features and investigate their effects on spheroid growth.

The ECM can also be characterised by its stiffness, rather than by its density and ribose concentration. ^27^ However, our focus was on understanding how ribose-induced collagen stiffening specifically affected cancer spheroid growth and invasion. As previously mentioned, ECM stiffness can be modulated through various methods, each impacting different properties of the ECM and ultimately influencing the behaviour of the cells, thereby affecting the spheroid growth and invasion. It would be interesting to explore how these different stiffening methods could be reproduced in the current model. Additionally, it has been observed that the timing of ECM stiffening can either inhibit or promote cancer cell invasion, highlighting the complex relationship between ECM stiffness and cancer cell invasion. ^19^ Matrix stiffening after cancer invasion begins promotes further spheroid invasion, while a stiff matrix surrounding the spheroid at the early stages prevents invasion.

In addition to integrating more ECM features, future iterations of our model could incorporate diffusible nutrients and cell death mechanisms. ^51,52^ Currently, our model employs an inhibition of proliferation function based on the number of neighbouring cells and ECM density (Equation

(9)). While this approach provides a basic setup, a more comprehensive model would directly account for the effect of nutrient availability and cell death on proliferation dynamics. However, incorporating these additional components into a hybrid model poses challenges, particularly with respect to parameter validation and mathematical function accuracy. This is worsened by the scarcity of relevant data. For instance, we lack the data necessary to distinguish between the proliferation and death of cancer cells within the spheroid.

It is also important to note that our model operates within a constrained 3-dimensional space. Although we conceptually address the impacts of the ECM on cell migration in 3D, all simulations and analyses have been conducted in 2D. This limitation may hinder our ability to fully capture the complexities of 3D environments and their effects on spheroid behaviour.

In conclusion, the interactions between cancer cells and the extracellular matrix in 3D cancer spheroid growth are intricate and not yet fully understood. Our proposed model represents an initial attempt to account for the chemical and mechanical interactions within this context, paving the way for future research that integrates additional ECM properties and environmental factors.

## Supporting information

invasive_rib0

non-invasive_rib0

## Acknowledgments

MB would like to give a special thanks to Kieran Atkins for sharing his programming expertise, which was crucial in cracking the code. We acknowledge the use of Grammarly (version 14.1200.0) and chatGPT-4 for text editing and to identify improvements in the writing style.

## Supplemental material

**Figure S1:**
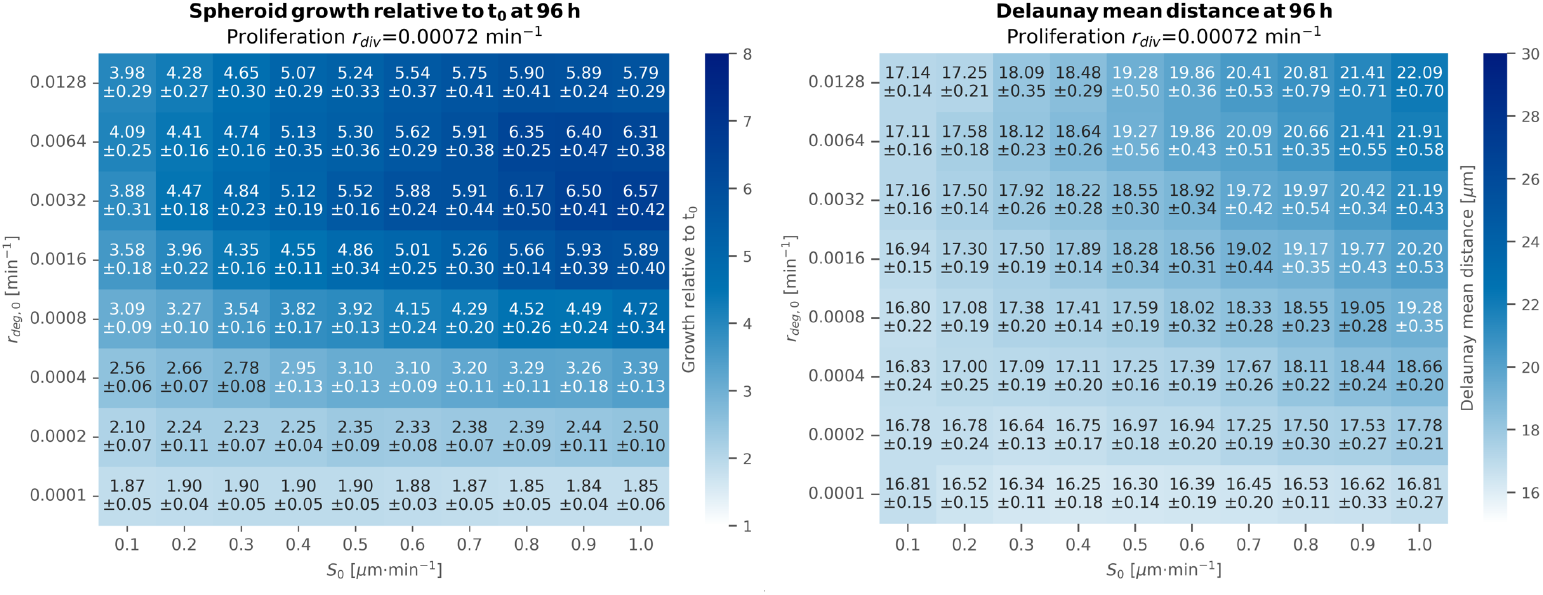
Heatmaps showing the effects of maximum cell-ECM interaction speed *S*_0_ (*x*-axis) and degradation rate *r*_*deg*,0_ (*y*-axis) on spheroid area growth relative to the initial time *t*_0_ (left) and Delaunay mean distance (right) after 96 h with ribose concentration 0 mM and proliferation rate *r*_*div*_ = 0.00072 min^−1^. Mean values and standard deviation over 10 replicates of the spheroid area growth relative to the initial time and Delaunay mean distance are shown. The colour intensity represents the mean values of the spheroid area growth relative to the initial time ranging from 1 to 8 and Delaunay mean distance ranging from 10 *µ*m to 30 *µ*m, as shown in the colour bars.

**Figure S2:**
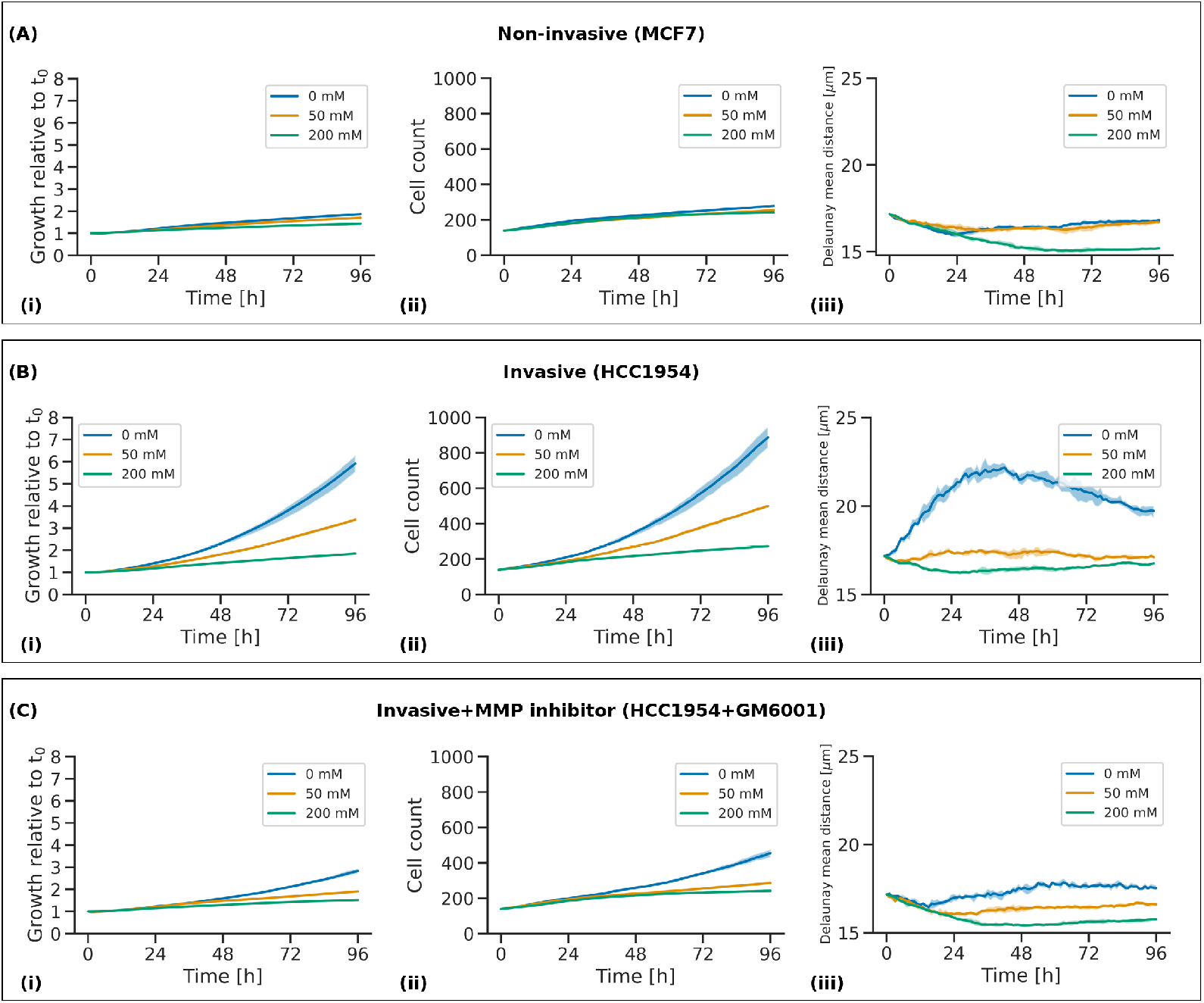
Results of simulations for non-invasive (MCF7) cells **(A)**, invasive (HCC1954) cells **(B)**, and invasive cells with the addition of MMP inhibition (HCC1954+GM6001) **(C)**) with ribose concentrations of 0 mM, 50 mM and 200 mM. Line plots of change over time of spheroid growth relative to the initial time *t*_0_ **(i)**, cell count **(ii)** and Delaunay mean distance **(iii)** are shown. The ribose concentrations are represented in blue for 0 mM, orange for 50 mM and green for 200 mM. Mean and 25th/75th percentile over 10 replicates are shown.

## Data availability statement

The datasets presented in this study can be found in online repositories. The names of the repository/repositories and accession number(s) can be found below: https://github.com/Margherita-Botticelli/PhysiCell-cancer-spheroid-ecm-stiffness.

